# Synaptic vesicle pools are a major hidden resting metabolic burden of nerve terminals

**DOI:** 10.1101/2020.11.16.385575

**Authors:** Camila Pulido, Timothy A. Ryan

## Abstract

The human brain is a uniquely vulnerable organ as interruption in fuel supply leads to acute cognitive impairment on rapid time scales. The reasons for this vulnerability are not well understood, but nerve terminals are likely loci of this vulnerability as they do not store sufficient ATP molecules and must synthesize them on-demand during activity or suffer acute degradation in performance. The requirements for on-demand ATP synthesis however depends in part on the magnitude of resting metabolic rates. We show here that, at rest, synaptic vesicle (SV) pools are a major source of presynaptic basal energy consumption. This basal metabolism arises from SV-resident V-ATPases compensating for a hidden resting H^+^ efflux from the SV lumen. We show that this steady-state H^+^ efflux is 1) mediated by vesicular neurotransmitter transporters, 2) independent of the SV cycle, 3) accounts for ~half of the resting synaptic energy consumption and 4) contributes to nerve terminal intolerance of fuel deprivation.

- Nerve terminals consume large amounts of ATP even at rest
- A large fraction of the resting metabolic burden arises from the V-ATPase
- Steady-state V-ATPase activity is driven by a synaptic vesicle H^+^ leak
- This H^+^ leak is mediated vesicular neurotransmitter transporters

## Introduction

The human brain generally has a very small safety factor with respect to fuel supply, such that when blood glucose levels drop by only ~ 2-fold, severe neurological consequences ensue. As a result, even brief interruptions in blood flow that restrict delivery of glucose & oxygen, key elements necessary for ATP synthesis, rapidly lead to severe neurological impairment. This metabolic vulnerability is heightened by the fact that the brain is also the most energetically expensive organ, consuming ~ 20% of the body’s fuel intake but representing only ~2.5% of the mass. A large fraction of energy consumption is driven by electrical activity in the brain, however two pieces of evidence support the idea that the brain tissue has a large resting metabolic burden decoupled from action potential (AP) firing: 1) even in a vegetative state where electrical activity is severely curtailed, cerebral glucose consumption decreases by only 2-fold (Laureys and Schiff, 2012; Levy et al., 1987); 2) deep anesthesia in rodents that lead to iso-electric EEGs (i.e. no detectable electrical activity) only reduces brain O_2_ consumption ~2-fold (Nemoto et al., 1996). The presence of large resting metabolic rates likely impose significant constraints on how well different neuronal compartments can adapt to changing metabolic needs. The molecular origin of this high resting metabolism however is not well understood, although it has generally been assumed that activity of the Na+/K+ ATPases, compensating for leak-currents across the plasma membrane, is the principal basal metabolic driver since this is also a principal energy consumer during activity (Attwell and Laughlin, 2001). Regardless of the molecular origin of basal metabolic rates, these processes constitute a constant drain on resources and will determine the ability of a given compartment to withstand temporary limitations in fuel supply, since any locally produced ATP is already “tapped’ for handling resting needs. We previously showed that nerve terminals are particularly vulnerable to fuel withdrawal, and rapidly fail to recycle SVs upon deprivation of a suitable combustible carbon source (Ashrafi et al., 2020; Ashrafi et al., 2017; Rangaraju et al., 2014). We wondered if this vulnerability might be exacerbated by local high basal metabolic rates and designed experiments to probe the magnitude of presynaptic basal energy consumption, its molecular origins and its impact on fuel intolerance. Here we show that nerve terminals have a high resting metabolic energy demand, independent of electrical activity and that synaptic vesicle (SV) pools are a major source of basal energy consumption in this compartment while the plasma membrane Na+/K+ ATPase is only a minor contributor. We demonstrate that this basal metabolism arises from SV-resident V-ATPases compensating for a previously unknown constant H^+^ efflux from the SV lumen. We show, 1) that this steady-state H^+^ efflux is mediated by the vesicular neurotransmitter transporter, independent of the SV cycle, 2) that it accounts for ~half of resting synaptic energy consumption and 3) that suppression of this transporter activity significantly improves the ability of nerve terminals to withstand fuel withdrawal. Our findings underscore why nerve terminals are susceptible to metabolic compromises, as in addition to regulating ATP production in responses to activity, they must constantly meet a large local metabolic burden.

## Results

### The vesicular proton pump, not the plasma membrane Na^+^/K^+^ ATPase, accounts for a large resting basal ATP consumption in nerve terminals

We developed a quantitative approach to determine the magnitude of the resting metabolic load at synapses. We measured the kinetics of presynaptic ATP (ATP_presyn_) decline in resting synapses of dissociated hippocampal neurons after acute fuel deprivation using a quantitative genetically-encoded optical reporter, Syn-ATP (Rangaraju et al., 2014) (Figure 1A). In the presence of the Na^+^ channel blocker tetrodotoxin (TTX), replacing glucose with the non-metabolizable 2-deoxyglucose (2DG) led to a rapid continuous depletion in ATP_presyn_ down to 32% ± 1.4 (n =13) of the original ATP_presyn_ levels in 25min (Figure 1B, gray) with an initial slope of ~ 5.4%/min. We previously showed that under resting conditions ATP_presyn_ is ~1.4 mM. This corresponds to an initial basal metabolic consumption of ~3100 ATP/s per nerve terminal (see methods). The action of the Na^+^/K^+^-ATPase to restore ionic gradients is considered to be one of the largest metabolic costs in the active brain (Attwell and Laughlin, 2001; Hallermann et al., 2012; Lennie, 2003). In the absence of AP firing, the Na^+^/K^+^-ATPase compensates for any leak currents across the plasma membrane. We sought to determine the extent to which this activity might contribute to the resting metabolic load by examining the kinetics of ATP depletion during fuel deprivation as above, but in the presence of ouabain, a potent inhibitor of the Na^+^/K^+^-ATPase pump. Surprisingly, inhibiting the Na^+^/K^+^-ATPase had no measurable impact on the kinetics of ATP depletion (Figure 1B, blue): in the presence of 1mM ouabain, after 25min of 2DG incubation, ATP_presyn_ decreased to very similar levels as controls (normalized depletion mean ± SEM; Ctrl: 1 ± 0.021 (n=13) vs ouabain: 0.95 ± 0.077 (n=14); Figure 1C). These experiments revealed a surprisingly low activity of Na^+^/K^+^-ATPase in the absence of AP firing and determined that this pump is not a significant contributor to the resting metabolic load at nerve terminals.

**Figure 1.**
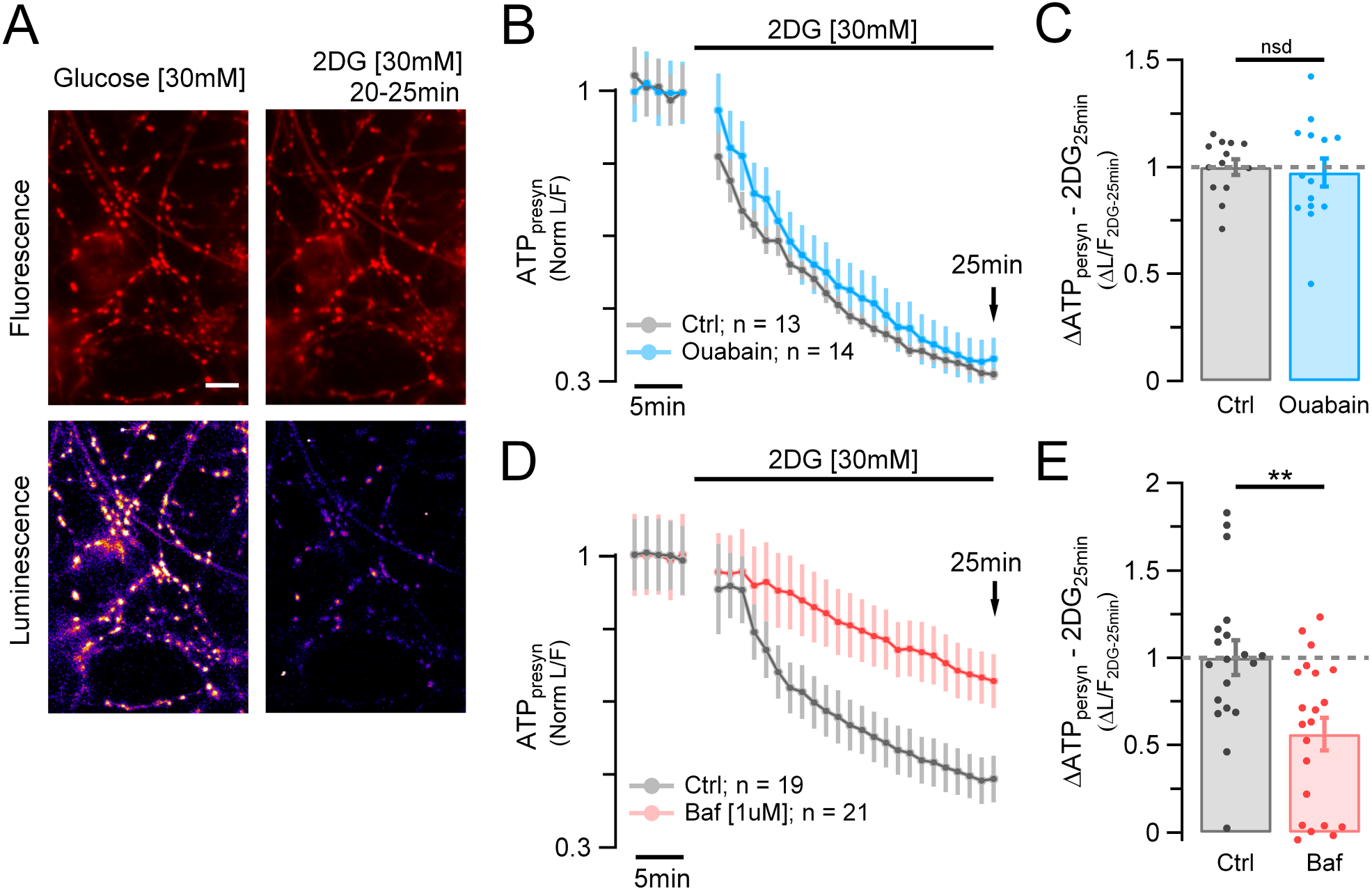
Synaptic vesicle V-ATPase, not the plasma membrane Na+/K+-ATPase, is a primary energy burden in resting synapses. (A) Syn-ATP fluorescence (F) (top) and luminescence (L) (bottom) images acquired from primary hippocampal neurons in glucose (left) and 25 min after replacing glucose with 2DG (right) in the presence of TTX (five frame average taken at 20-25 min). Scale bar, 10um. (B and D) Average ATP_presyn_ (L/F intensity ratio) time course normalized to baseline measured in glucose. (B) Ensemble average time course from neurons incubated in 2DG (n = 13; gray trace) or incubated in 2DG + ouabain [1mM] (n =14; blue trace). (C) Average of ΔATP_presyn_ (ΔL/F) values after 25min in 2DG (2DG_25min_) in the presence of ouabain (blue dots) normalized to control (gray dots): mean ± SEM: 0.95 ± 0.077 vs 1 ± 0.021 (nsd). (D) Ensemble average time course from neurons incubated in 2DG (n = 19; gray trace) or incubated in 2DG + bafilomycin [1uM] (n = 21; red trace). (E) Average ΔATP_presyn_ (ΔL/F) values after 25min in 2DG (2DG_25min_) in the presence of bafilomycin (red dots) normalized to control (gray dots): mean ± SEM: 0.56 ± 0.093 vs 1 ± 0.1 (p < 0.002).

We therefore considered alternative mechanisms at presynaptic terminals to explain basal metabolic dynamics. SVs utilize the proton motive force generated by vesicle-resident V-ATPases, with 1-2 copies per vesicle (Takamori et al., 2006), each hydrolyzing three ATP molecules per ten protons translocated (Abbas et al., 2020) to power neurotransmitter uptake into the vesicle lumen (Eriksen et al., 2020). Although one expects ATP expenditure for filling vesicles with neurotransmitter, it is less clear in a resting vesicle pool if there is any residual activity that might account for part of the resting metabolic load. We tested for this possibility by examining basal ATP consumption (in TTX and 2DG) in the presence of bafilomycin, a potent V-ATPase inhibitor. These experiments showed that, unlike with blockade of the plasma-membrane Na^+^/K^+^ pump, blocking V-ATPases significantly reduced basal ATP consumption. The initial presynaptic ATP depletion rate in in the presence of 1μM bafilomycin was ~3 times slower that in control (t1/4-baf = 22.01min ± 3.6 (n = 21), t1/4-ctrl = 7.69min ± 1.36 (n = 19); p < 0.001; Figure 1C), leaving ~69.91% ± 6.5 of the original ATP at 25 min compared to just a 46.41% ± 5.6 in controls (Figure 1D; p < 0.002). The ATP values normalized to Ctrl_25min_ are, respectively, 0.56 ± 0.093 vs 1 ± 0.1 (p < 0.002; Figure 1E). These data unmask the V-ATPase as a main energy burden in resting synapses that accounts for ~half of the resting ATP consumption, i.e. ~1550 ATP/s per nerve terminal. Given the large total number of nerve terminals in the brain, and that this burden will be present constantly, these results imply the resting SV pools in total likely constitute a significant energy burden in the brain.

### A “hidden” H^+^ flux from the SV lumen is present even in the absence of SV exocytosis and recycling

V-ATPases are electrogenic pumps and as such only stop burning ATP when the energy barrier for moving a proton against a chemical potential exceeds the energy provided by hydrolyzing ATP. Our results imply that the H^+^ gradient in SVs is constantly dissipated but restored by the V-ATPase, which when blocked would lead to alkalization of SVs. To test this hypothesis, we measured changes in SV luminal pH (pH_Lum_) from resting nerve terminals of neurons expressing vGlut-pHluorin (vG-pH) before and after blocking the V-ATPase with bafilomycin (Sankaranarayanan and Ryan, 2001). Prior to blocking the proton pump, the basal fluorescence of boutons expressing vG-pH remained stable over time (2 min baseline measurements, Figure 2B). However, upon addition of bafilomycin, the vG-pH fluorescence immediately began to increase at a rate of 0.117% ± 0.01%F_max_/s (n = 13), where F_max_ is the fluorescence at pH of 6.9, the pH of the cytoplasm, referred to hereon as F_pH6.9_ (Figure 2B and C; black trace). These data indicate that there is a steady-state efflux of H^+^ from SVs, which in turn is being compensated by the action of the V-ATPase, at the expense of ATP hydrolysis. This H^+^ efflux showed a reasonably strong temperature dependence, as lowering the temperature from 37°C to 25°C reduced the rate by a factor of ~2 (Figure S1). Similar results were obtained using an alternate V-ATPase blocker, folimycin (Figure S2) with only minor differences between the potency of the two pump blockers, which saturate near ~500nM. These data imply that the flux of H^+^ is occurring from resting SVs, however such changes in pH would also occur if the SVs underwent spontaneous exocytosis, releasing the protons to the extracellular space. To determine the possible contribution of this mechanism to our measurements we carried out two types of experiments. The first was to repeat the H^+^ flux measurements in synapses where we strongly suppressed exocytic mechanisms by either genetically expressing tetanus-toxin light chain (TeNT) which cleaves the major vSNARE for synaptic exocytosis, VAMP-2 (Gaisano et al., 1994; Schiavo et al., 1992), or by suppressing expression of Munc13-1/2 using an shRNA targeting the genes encoding these proteins (Calloway et al., 2015). In hippocampal neurons, a combination of Munc13-1 & 2 is required for all known forms of synaptic vesicle exocytosis (Varoqueaux et al., 2002). We verified that either treatment prevented any AP driven exocytosis of vG-pH (Figure 2A) and then examined the time course of vG-pH fluorescence change following application of bafilomycin. These experiments show conclusively that the fluorescence increase following proton pump block occurs without any exocytosis: Ctrl: 0.117% F_pH6.9_/s ± 0.001 vs Munc13-KD: 0.124% F_pH6.9_/s ± 0.018 (n = 6) and TeNT: 0.119% F_pH6.9_/s ± 0.016 (n = 6); Figure 2B and C).

**Figure 2.**
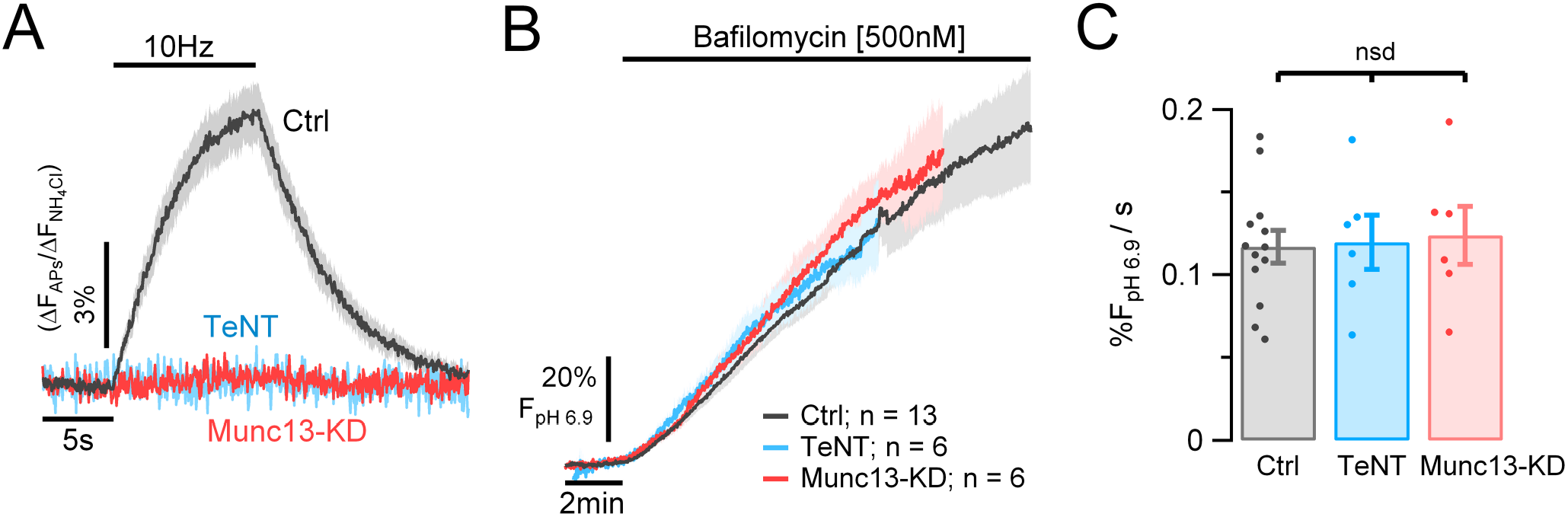
Resting synaptic vesicles have a constant H^+^ efflux. H^+^ flux from synaptic vesicles measured in hippocampal neurons expressing vG-pH. (A) Average vG-pH traces in response to 100APs (10Hz) in control neurons (Ctrl, gray trace, n = 13) or in neurons where exocytosis is genetically suppressed by either ablating expression of Munc13 (Munc13-KD, red trace, n = 6) or expressing tetanus toxin light chain (TeNT, blue trace, n = 6). ΔF values are normalized to maximal ΔF from NH_4_Cl treatment (ΔF/ΔF_NH4Cl_). (B) vG-pH average traces of Ctrl, Munc13-KD and TeNT neurons measured in presence of TTX before and after perfusion with bafilomycin. Fluorescence expressed as a percentage of the fluorescence expected for pH = 6.9 (%F_pH6.9_) based on perfusion with NH_4_Cl (see methods). ± SEM intervals are indicated by shaded colored areas. (C) Bafilomycin application causes SV vG-pH fluorescence to increase at the same rate in all three conditions, and thus is not related to exocytosis. Average rates of alkalization for Ctrl, Munc13-KD and TeNT neurons measured over the first 6 min in bafilomycin, respectively as: 0.117% F_pH6.9_/s ± 0.001, 0.124% F_pH6.9_/s ± 0.018 and 0.119% F_pH6.9_/s ± 0.016.

A second approach was to examine the apparent H^+^ flux in synapses that show no evoked synaptic responses even in their native state. Previously, we showed that for any given axon, a small portion (~15-20%) of synaptic boutons show no evoked exocytosis in response to AP bursts and are intermixed randomly with secretion-competent boutons in the same axon (Kim and Ryan, 2010). We first measured the size of action potential evoked signals following a 100AP-10Hz stimulus train across all boutons using vG-pH normalized to the maximal signal obtained during NH_4_Cl perfusion (Figure 3A left and B). We defined a “silent” synapse as one that failed to show an AP-evoked change greater than 1.2 standard deviation (SD) of the pre-stimulus baseline (Figure 3A, right) and “active” synapses as those whose response cross this threshold (Figure 3A, middle). Using these criteria, 80% ± 4.7 (n = 13) of synaptic boutons in our experiments were scored as active (Figure 3C), consistent with previous findings (Cazares et al., 2016; Kim and Ryan, 2013). Subsequently, we carried out proton-pump blocking experiments in the same neurons in TTX. We then compared the H^+^ efflux rates in active and silent synapses. These experiments showed that H^+^ efflux rates from individual boutons were unrelated to synaptic state defined as above (Silent: 0.133% F_6.9pH_/s ± 0.021 vs active: 0.116% F_6.9pH_/s ± 0.013; Figure 3D and E), solidifying the notion that resting synaptic vesicle pools have substantial sustained V-ATPase activity at the expense of ATP consumption.

**Figure 3.**
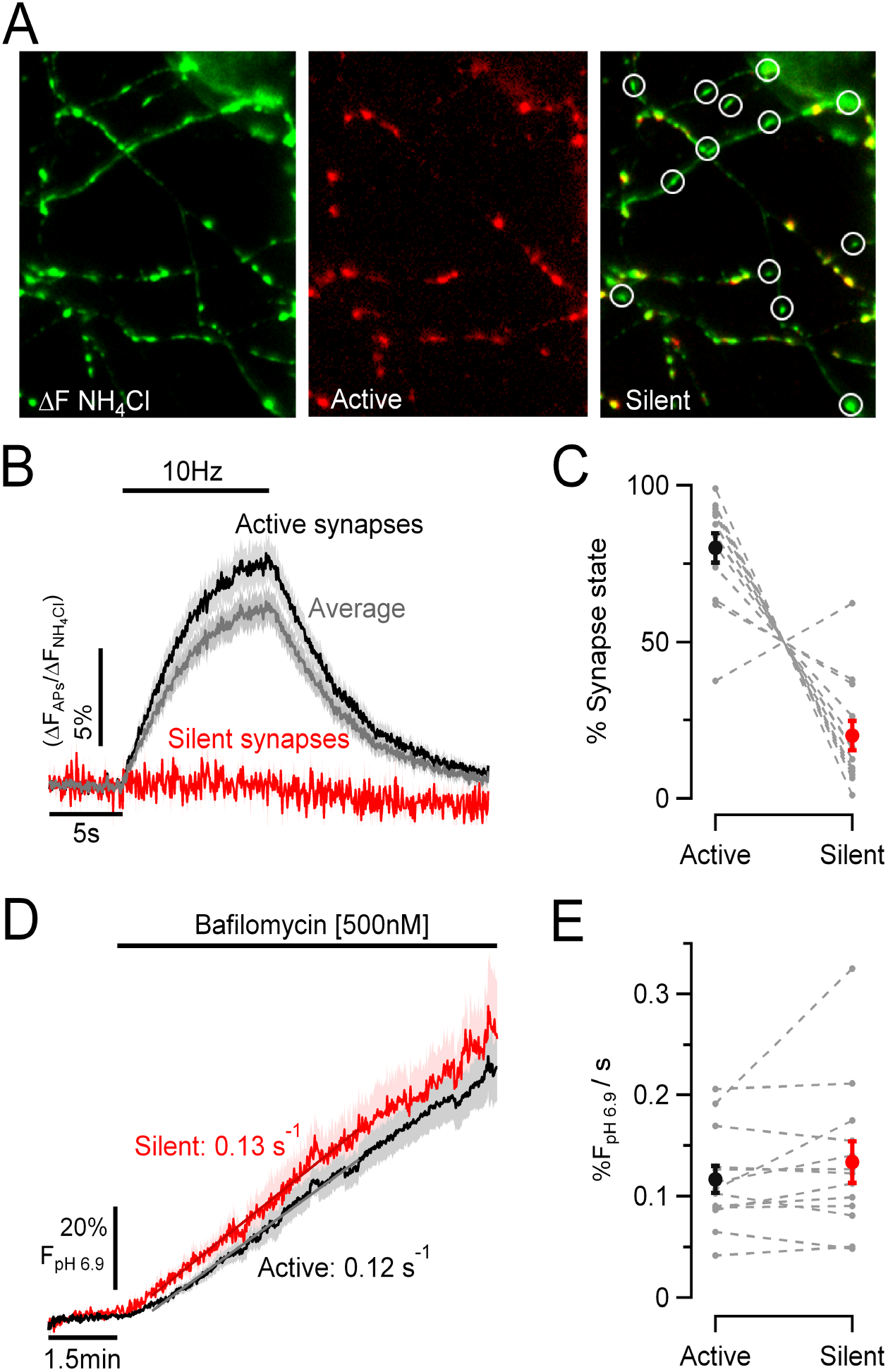
Synaptic vesicle H+ efflux is not correlated with individual synapse release properties. (A) Representative images showing variation in synaptic release in an individual neuron expressing vG-pH. Perfusion with NH_4_Cl perfusion (left; pseudo-colored green) reveals expression pattern of vG-pH across synapses. vG-pH responses to electrical activity (100APs/10Hz, middle; pseudo-colored red) shows that a portion of the nerve terminals are silent (right; overlay between left and middle images) indicated by circles. (B) Average vG-pH response to stimulation from active (black), silent (red) and total synapses (gray) per neuron (n =13); where silent synapse is defined as showing responses less than 1.2 SD of the pre-stimulus baseline. (C) Percentage of nerve terminals in each category on a neuron-by-neuron basis (dashed gray lines) shows that ~ 80% of nerve terminals show robust responses and 20% are silent. (D) Ensemble average SV H+ efflux kinetics measured in TTX and bafilomycin for active and silent synapses across all neurons (n=13) or on a neuron-by-neuron basis. (E) show that the SV H+ efflux is unrelated to individual bouton exocytic properties (active: 0.116%F_pH6.9_/s ± 0.013 and silent: 0.133% F_pH6.9_/s ± 0.021). Individual rates are shown in dashed gray lines (n = 13).

### vGlut mediates the steady-state H^+^ flux from synaptic vesicles

We next turned to identifying the molecular basis of the SV H^+^ efflux. All SV types utilize vesicular neurotransmitter transporters driven in part by the energy provided by the V-ATPase to drive the filling process via an alternating access mechanism (Forrest et al., 2008). Although some vesicular neurotransmitter transporters are known to exchange H^+^ with neurotransmitter during the pumping process, it has been difficult to ascertain this for vGlut family members (Eriksen et al., 2020). We wondered if vGlut itself might mediate the proton “leak”. In order to test this hypothesis, we used shRNA-mediated knock-down of vGlut1 expression (Pan et al., 2015) in neurons expressing a pHluorin-tagged synaptophysin (Syphy-pH). Consistent with our previous findings (Pan et al., 2015), loss of vGlut1 did not impair exocytosis (Figure 4A, inset). Measurements of H^+^ efflux (in bafilomycin) from the SV pool in control neurons expressing Syphy-pH were similar to those measured with vG-pH (0.12% F_pH6.9_/s ±0.016, n = 12; Figure 2C and 4B), however, knock-down of vGlut1 significantly reduced the H^+^ efflux rate (vGlut1-KD: 0.052% F_pH6.9_/s ± 0.006 (n =11), p < 0.0005; Figure 4A and B). This reduction in SV H^+^ efflux in vGlut1 KD neurons was fully reversed upon expression of the human variant of vGlut1, which was resistant to the shRNA used for rat (vGlut1-rsc: 0.12% F_pH6.9_/s ± 0.022 (n = 6), p < 0.5 Figure 4A and B). Although we previously showed that this shRNA-mediated ablation suppresses vGlut1 expression by ~93% (Pan et al., 2015), it is possible the remaining H^+^ efflux following knockdown arises from residual vGlut1. This transporter is thus necessary for H^+^ efflux in resting SVs.

**Figure 4.**
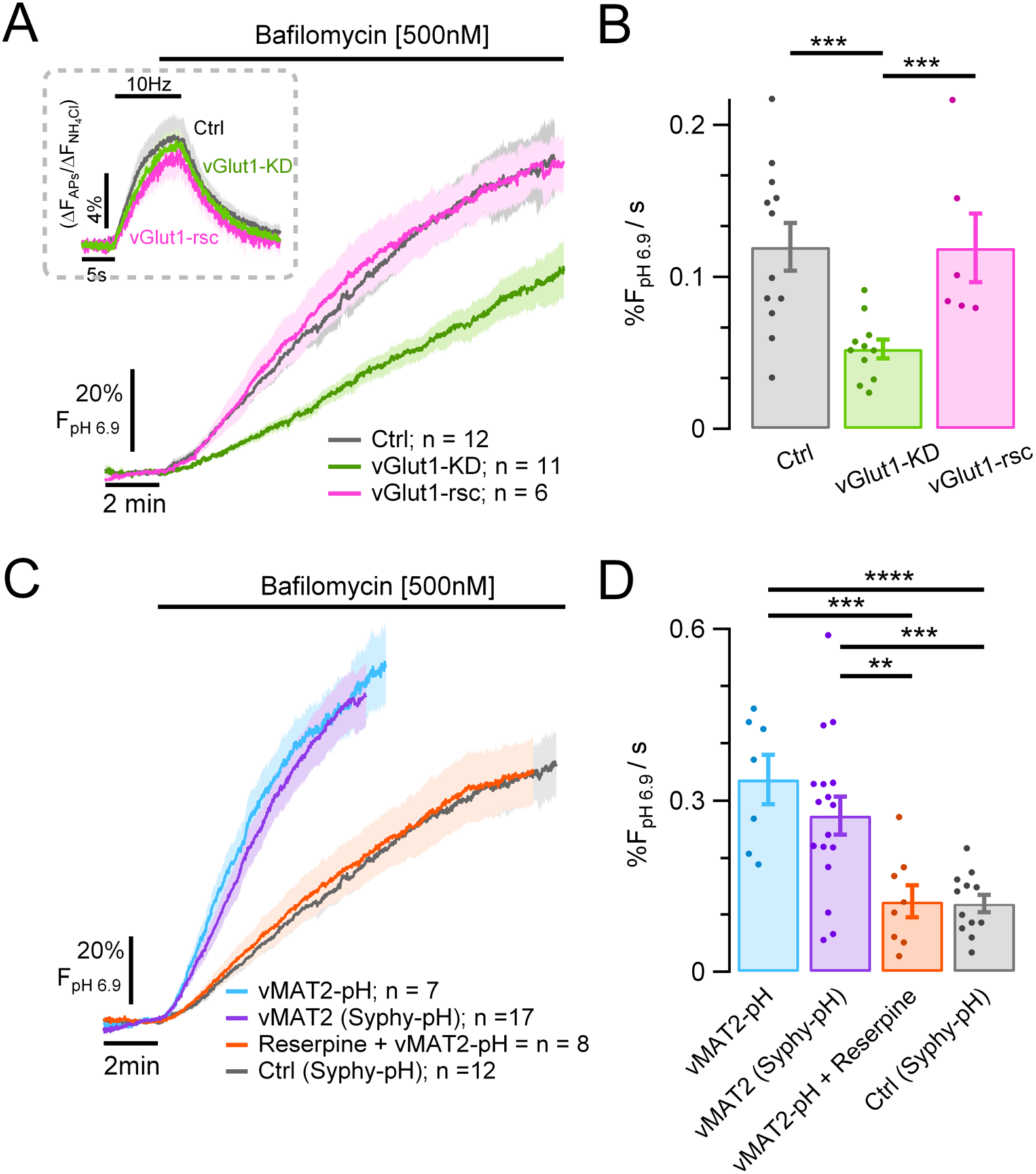
Vesicular neurotransmitter transporters (i.e., vGlut1 and vMAT2) mediate a large fraction of the resting SV H^+^ efflux. (A and B) SV H^+^ efflux was measured in the presence of TTX and bafilomycin in hippocampal neurons expressing either Syphy-pH (control, gray trace), Syphy-pH and an shRNA suppressing expression of vGlut1 (vGlut1-KD, green trace) or Syphy-pH, vGlut1-KD and a human variant of vGlut1, resistant to the ShRNA (vGlut1-rsc, pink trace). (A) Control, vGlut1-KD and vGlut1-rsc show similar exocytic responses to electrical stimulation (inset), but vGlut1-KD neurons show much lower SV H^+^ efflux. (B) SV H^+^ efflux rates in Ctrl: 0.12 %F_pH6.9_/s ± 0.016 (n = 12) vs vGlut1-KD: 0.052%F_pH6.9_/s ± 0.0062 (n = 11), p < 0.0005. SV H^+^ efflux is fully recovered in neurons where vGlut1 was rescued (0.12%F_pH6.9_/s ± 0.022 (n = 6) vs vGlut1-KD p < 0.0005). (C) Hippocampal neuronal expression of the exogenous transporter vMAT2, either coupled to pHluorin (vMAT2-pH, blue trace, n = 7) or co-expressed with Syphy-pH (vMAT2, violet trace, n = 17) led to a faster SV H^+^ efflux in presence of bafilomycin, compared with control neurons expressing native vGlut1 (Syphy-pH, gray trace, n = 12). This increase in H^+^ flux is completely abolished by application of 100nM reserpine (red trace, n = 8). (D) Bar plot of the SV H+ efflux rates of vMAT2-pH (0.34%F_pH6.9_/s ± 0.043), vMAT2 (0.28%F_pH6.9_/s ± 0.033), reserpine (0.124%F_pH6.9_/s ± 0.029) and control (0.12%F_pH6.9_/s ± 0.016). vMAT2-pH vs: Ctrl, p < 0.0001 and reserpine, p < 0.001. vMAT2 vs: Ctrl, p < 0.0003 and reserpine, p < 0.002.

### Exogenously expressed vMAT2 drives a reversible SV H+ efflux in hippocampal neurons

Although our data show that vGlut is necessary for a large fraction of the resting H^+^ flux from SVs, whether vGlut is a direct or indirect mediator of H^+^ flux is not clear from our experiments. It is possible that other proteins responsible for maintaining SV ionic flux balances during glutamate uptake are responsible for the H^+^ current. To address this problem, we sought to determine if other vesicular neurotransmitter transporters, in particular ones considered to be *bona fide* proton exchangers, could mediate a H^+^ leak from SVs. We chose to measure H^+^ fluxes in neurons where we expressed an exogenous neurotransmitter transporter for which there is no cognate neurotransmitter molecule, and which additionally can be blocked acutely using a pharmacological inhibitor. As hippocampal neurons do not express tyrosine hydroxylase (TH), they do not synthesize dopamine. Dopamine and serotonin are packaged into SVs in neurons that express TH by vMAT family members. Additionally, vMATs are known to exchange 2 protons for each monoamine taken up in the dopaminergic SV lumen (Johnson et al., 1981; Knoth et al., 1981). Expression of vMAT2-pHluorin (vMAT2-pH) (Onoa et al., 2010; Pan and Ryan, 2012) in hippocampal neurons was localized to nerve terminals with very low surface expression (not shown). Similar to our measurements with either vG-pH or Syphy-pH, application of bafilomycin led to a prompt increase in vMAT2-pH fluorescence. The rate of increase was ~ 2.8-fold greater than in nerve terminals expressing Syphy-pH (i.e. without vMAT2; vMAT2-pH: 0.34% F_pH6.9_/s ± 0.043 (n = 7) vs Syphy-pH: 0.12% F_pH6.9_/s ± 0.016 (n = 12); p < 0.0001; Figure 4C and D). As with vGlut1, the H^+^ flux was independent of tagging with pHluorin as expression of vMAT2 along with Syphy-pH showed very similar H^+^ flux rates compared to vMAT2-pH neurons (0.28% F_pH6.9_/s ± 0.033, n = 17; p < 0.2). Importantly however, application of reserpine, a potent inhibitor of vMAT2, completely suppressed the increase in H^+^ flux (0.124% F_pH6.9_/s ± 0.03, n = 8; p < 0.001). These data, together with the vGlut1 KD experiments, strongly support the idea that the transporters themselves can mediate the SV H^+^ efflux.

### vGlut1-mediated SV H^+^ accounts for ~50% of the high resting ATP consumption

Our data shows that a constant proton flux accounts for ~50% of resting synaptic ATP consumption (Figure 1D and E), and that this in turn arises because of a constant H^+^ efflux mediated by vGlut1. These data make a strong prediction that loss of vGlut1 should decrease resting ATP consumption, similar to simply blocking the V-ATPase. To test this hypothesis, we measured basal ATP consumption rates (in TTX and 2DG) in neurons where vGlut1 expression was suppressed. After removal of vGlut1, the ATP (expressed as L/F) rate of decay in presence of 2DG was slower than control (t1/4:16.8min ± 1.65 (n = 12) vs 7.69min ± 1.36 (n = 19); p < 0.0002; Figure 5A); after 25min of incubation 63.03%± 5.1 remained from the original ATP content, which is significantly higher than the 46.41% ± 5.6 left in control neurons (p < 0.01). Figure 5B shows normalized ΔATP_presyn_ values of vGlut1-KD with respect to Ctrl_25min_ (0.67 ± 0.056 vs 1 ± 0.1, p < 0.01). This reduction in the basal ATP consumption in vGlut1-KD neurons was fully recovered upon expression of vGlut1-rsc (values normalized to Ctrl_25min_: 1.044 ± 0.052 (n = 7), p < 0.7). Furthermore, ATP_25min_ percentage in vGlut1-KD neurons was similar to the ones obtained with bafilomycin (values normalized to Ctrl_25min_: bafilomycin: 0.56 ± 0.093 (n = 21) vs vGlut1-KD: 0.67 ± 0. 056 (n = 12); p < 0.7; Figure 1E vs 5B). Overall, the results reveal that the synaptic vesicle pool is a major source of metabolic activity even in the absence of electrical activity and neurotransmission.

**Figure 5.**
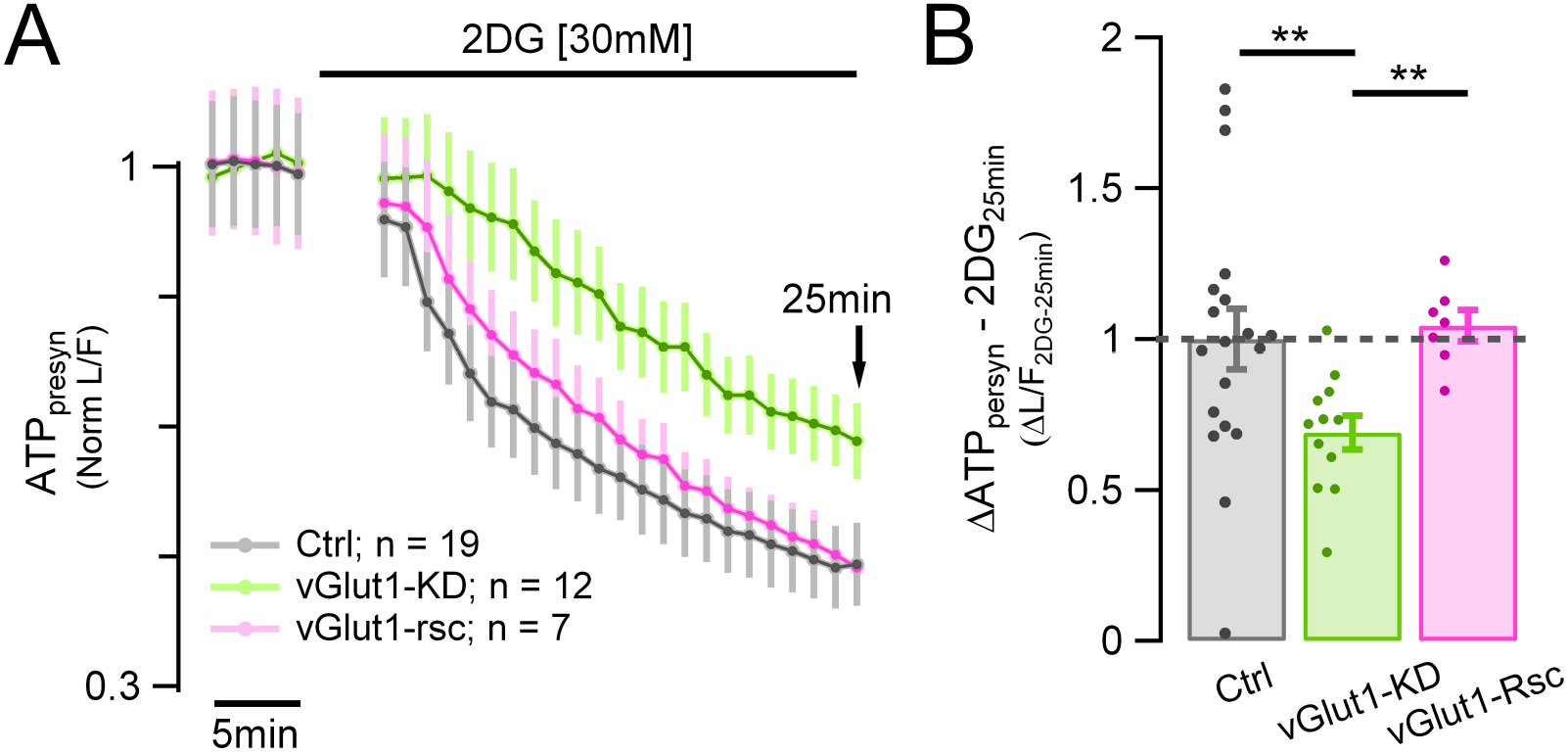
The SV H^+^ efflux mediated by vGlut1 is compensated by the action of the v-ATPase. (A) The resting ATP_presyn_ consumption measured using Syn-ATP in TTX and 2DG expressed as normalized L/F in vGlut1-KD (green trace; n = 12), vGlut1-KD + vGlut1-rsc (pink trace; n = 7) and control neurons (gray trace; n = 19). (B) Suppression of vGlut1 expression lowers the resting ATP consumption by 33% (measured after 25 min in 2DG, 2DG_25min_) in vGlut1-KD neurons (green dots) normalized to control neurons (gray dots): mean ± SEM: 0.67 ± 0.056 vs 1 ± 0.1, (p < 0.01). Resting ATP_presyn_ consumption is fully rescued compared to Ctrl levels (vGlut1-rsc: mean ± SEM: 1.05 ± 0.052, p < 0.7).

### Suppressing vGlut1 expression sustains nerve terminal function in restricted fuel conditions

Our data imply that the H^+^ efflux from SVs contributes to a high resting metabolic rate. As one of the primary manifestations of a compromise in fuel availability at nerve terminals is an arrest in SV recycling, we speculated that suppressing vGlut1 expression would allow nerve terminals to sustain SV recycling for longer periods when the fuel supply is removed, similar to how improving fuel efficiency in would extend the driving range of a vehicle. To test this idea, we used Syphy-pH expressed in hippocampal neurons to measure SV recycling kinetics in response to activity in sequential rounds of 50 AP 10 Hz bursts delivered at 1 min intervals immediately after all glucose was removed in both control neurons and those in expressing an shRNA targeting vGlut1 (Figure 6). Under these conditions, all energetic needs must presumably be met by either residual ATP or intermediates in ATP production since no external combustible carbon source is being provided. In control neurons, removal of glucose led to the gradual slowing of SV recycling, here shown color-coded with respect to stimulus round (Figure 6A), with the onset of slowing apparent in 5 minutes and a complete arrest apparent at the sixth round of AP bursts (Figure 6A, C and D), consistent with our previous findings (Ashrafi et al., 2017; Rangaraju et al., 2014). To express the impact of SV recycling quantitatively, we measured the fraction of the exocytic signal that remained a after defined post-stimulus time period (3 times constants of the decay in the first round), which we term the percentage of endocytic block (=~5% in the first round by definition for a perfect exponential decay). For a given neuron, we determined the number of stimulus rounds it took for SV recycling to fail to retrieve 50% of the exocytic signal at this time period, which in control neurons was on average 5.9 rounds ± 0.5 (n = 12; Figure 6D). In contrast when vGlut1 expression is suppressed, in the absence of any fuel, nerve terminals can now sustain typically 8.31 rounds ± 0.67 (n = 13) of recycling before SV recycling arrests (p < 0.006; Figure 6B, C and D). These experiments demonstrate that the energetic burden created by the H^+^ efflux mediated by vGlut1, strongly impacts the efficacy of the synaptic vesicle cycle when fuel availability is restricted.

**Figure 6.**
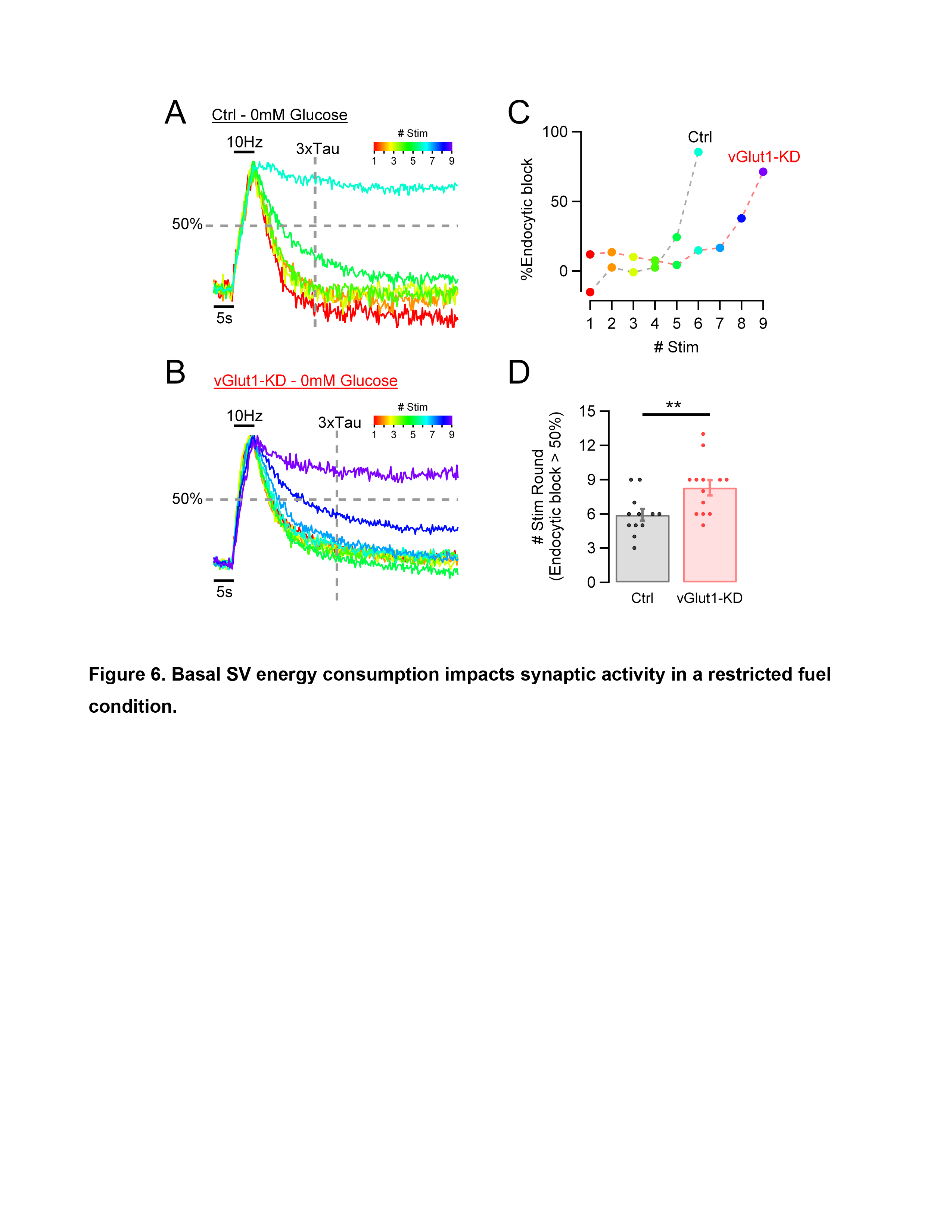
Basal SV energy consumption impacts synaptic activity in a restricted fuel condition. (A and B) synaptic activity measured in hippocampal neurons expressing Syphy-pH in the absence of glucose (0mM Glucose). Color-code traces represent responses to one stimulus round (50AP −10Hz) applied every minute (from red = 1^st^ round to violet = 9^th^ round). Responses are normalized to the peak. Vertical dashed line represents 3 times Tau measured at the first round. Horizontal dashed line represents 50% retrieval of the exocytic signal. (A) Traces to six stimulus rounds in a control neuron. (B) Traces to 9 stimulus rounds in a neuron suppressing expression of vGlut1 (vGlut1-KD). (C) percentage of endocytic block from the examples shown in panels a and b, measured each round at 3xTau time point. vGlut1 suppression prolongs synaptic activity by 3 rounds compared to control neuron. (D) Number of rounds of stimulation before endocytic block exceeds 50% is smaller in neurons expressing vGlut1 (gray dots, n = 12) than in vGlut1-KD neurons (red dots, n = 13): mean ± SEM: 5.92 rounds ± 0.52 vs 8.31 rounds ± 0.67; p < 0.005.

## Discussion

Our studies provide a compelling explanation for the reason nerve terminals are so sensitive to metabolic compromise, and in turn potentially speaks to why brain tissue in general has a resting metabolic rate that is much higher than other tissues. The presence of synaptic vesicles themselves and their intrinsic reliance on the V-ATPase to power neurotransmitter filling have rendered them sensitive to routes of efflux for H^+^ from the vesicle lumen, which in turn sustains V-ATPase activity creating a constant energetic burden. Our experiments do not pin down the precise route of H^+^ efflux in glutamatergic vesicles but do show that vGlut is a critical mediator. Previous experiments (Eriksen et al., 2016), provided compelling evidence that luminal protons allosterically modulate glutamate transport by vGluts, an idea bolstered by recent ultrastructural studies of this transporter (Li et al., 2020). The recent structural insights argue for an exchange for Cl^-^ ions with glutamate during the transport process, that is facilitated by low luminal SV pH. In this model, the pH gradient created by V-ATPases is only dissipated during exocytosis, not by the transport process. Our data indicate that even in the absence of exocytosis, SVs dissipate the H^+^ gradient over a ~10-minute period in a manner that clearly depends on vGlut. This dissipation occurs in SV vesicles that have almost certainly reached a steady-state with respect to neurotransmitter filling, as it occurs in synapses that have either very low release probability (Figure 3), or in which exocytosis has been prevented genetically (Figure 2). Thus, it is unlikely that the H^+^ efflux requires active transport of glutamate into the vesicle. Expression of vMAT2 in a neuron that lacks monoamines also resulted in H^+^ efflux that was acutely prevented by a blocker of the transport mechanism. These data support the idea that the transporters themselves likely directly mediate H^+^ efflux but without neurotransmitter transport. A parsimonious explanation is that the energy barrier for switching between conformations in the vesicular neurotransmitter transporters can be driven at a low rate by thermal fluctuations, even in the absence of cargo (neurotransmitter) binding on the cytosolic facing conformation. If this occurs and if protons are normally exchanged for neurotransmitter as part of the alternate access model, a proton would then get ejected from the SV vesicle lumen, essentially driven purely by thermal fluctuations.

It is interesting to note that expression of vMAT2 led to an almost 3-fold greater H^+^ efflux than that in native glutamatergic nerve terminals (Figure 4). Given that expression of pHluorin-tagged SV proteins leads to only one or two copies of tagged proteins per SV (Balaji and Ryan, 2007; Sinha et al., 2011), and that SV are estimated to have between 10-20 vesicular neurotransmitter transporters (Takamori et al., 2006) these data imply vMAT2 might have an intrinsically higher thermally-driven H^+^ efflux rate than vGlut1. Part of this might arise from a higher stoichiometry of H^+^/dopamine exchange compared to a putative H+/glutamate exchange, but the fact that expression of only one or two copies of vMAT2 transporter can lead to three-times greater efflux than all of the native vGlut1 in a glutamatergic terminal suggests that vMAT2 is intrinsically “leakier” to H^+^ than vGlut1. One could therefore predict that in dopaminergic terminals, where vMAT2 is natively expressed, the resting SV H^+^ efflux would be much larger. This in turn predicts an increased energetic burden at rest, perhaps making dopaminergic neurons inherently more susceptible to metabolic compromise.

These data have profound implications with respect to how energy balances in the brain are achieved and whether different neuronal populations might be more vulnerable than others to metabolic compromise due to the total load created by these SV pools. An important element of this is understanding how different neurons adjust ATP production. We previously showed that nerve terminals make use of both feedback and feedforward regulatory mechanisms for ATP production. Feedback mechanisms respond to increased ATP consumption and the ensuing changes in the AMP/ATP ratio (Ashrafi et al., 2017). This type of pathway would be well suited to adjust ATP levels as basal activity-independent change. Activity-dependent feed-forward mechanisms to modulate ATP production such as proposed for the lactate shuttle (Pellerin et al., 1998) or Ca^2+^-driven mitochondrial regulation (Ashrafi et al., 2020) might not compensate for large resting metabolic loads since they are inherently uncoupled from activity. Thus, depending on what regulatory strategies are available, different neurons may be differentially impacted by such high metabolic burdens. Although at face value using a vesicle filling method that results in a continuous ATP consumption appears inefficient, it may simply result from optimizing the speed with which SVs are filled with neurotransmitter. Faster transport is generally achieved by lowering the energy barrier for the key structural transition, which in turn implies a higher probability of occurring from simple thermal fluctuations. Given the vast number of synapses in the human brain, and the presence of hundreds of SVs at each these nerve terminals, this hidden metabolic cost of quickly returning synapses in a “ready” state comes at the cost of major ATP and fuel expenditure, likely contributing significantly to the brain’s metabolic demands and metabolic vulnerability.

## Methods

### Animals

All animal-related experiments were performed in accordance with protocols approved by the Weill Cornell Medicine IACUC. Wild-type rats were of the Sprague-Dawley strain (Charles River Strain code: 400, RRID: RGD_734476).

### Primary neuronal culture

Hippocampal CA1-CA3 neurons were isolated from 1 to 3 days old rats of mixed gender and plated on poly-ornithine-coated coverslips as previously described (Ryan, 1999). Calcium phosphate-mediated gene transfer was used to transfect 6–8-day-old cultures as described previously (Sankaranarayanan and Ryan, 2000) Neurons were maintained in culture media composed of MEM (Thermo Fisher Scientific S1200038), 0.6% glucose, 0.1 g/l bovine transferrin (Millipore 616420), 0.25 g/l insulin, 0.3 g/l glutamine, 5%–10% fetal bovine serum (Atlanta Biologicals S11510), 2% B-27 (Thermo Fisher Scientific 17504-044), and 4 mm cytosine b-d-arabinofuranoside. Cultures were incubated at 37°C in a 95% air/5% CO_2_ humidified incubator for 14–21 days prior to use.

### Plasmids Constructs

The following previously published DNA constructs were used: vGLUT1-pHluorin (Voglmaier et al., 2006), Munc13-1/2 shRNA and Syn-ATP (Rangaraju et al., 2014), vGlut1-shRNA and Syphy-pH (Pan et al., 2015), vMAT2-pH (Pan and Ryan, 2012), cyto-pH (gift of M.E. Hatten) and TeNT-LC (gift of M. Dong). DNA constructs made for this project: Human vGlut1 shRNA resistant rescue and vMAT2.

### Live cell imaging

Live-cell imaging was performed using a custom-built laser illuminated epifluorescence microscope with an Andor iXon+ (model #DU-897E-BV) back-illuminated electron-multiplying charge-coupled device camera that was selected for low dark currents. TTL-controlled Coherent OBIS 488 nm and 561 nm lasers were used for illumination. Images were acquired through a 40x 1.3 NA Fluar Zeiss objective. Experiments were performed at a clamped temperature of 37°C (or 25°C, Extended Data Figure 1) using a custom-built objective heater under feedback control. Action potentials were evoked by passing 1-ms current pulses, yielding fields of approximately 10 V cm-1 via platinum-iridium electrodes. Neurons were continuously perfused at 0.1 ml min-1 with a Tyrode’s solution containing (in mM) 119 NaCl, 2.5 KCl, 2 CaCl2, 2 MgCl2, 30 glucose, 0.01 6-cyano-7-nitroquinoxaline-2,3-dione (CNQX), 0.05 D, L-2-amino-5-phospho-novaleric acid (AP5) and 0.3 Tetradotoxin (TTX), buffered to pH 7.4 using 25mM HEPES. Tyrode’s solution without TTX was used for electrical stimulation experiments. Bafilomycin solutions were prepared freshly from a 250μM stock into a Tyrode’s + TTX solution and administrated as indicated in each experiment. In experiments without glucose, Tyrode’s molarity was compensated with either deoxy-glucose (2DG) or HEPES (0-Glucose).

### ATP Measurements

Luminescence imaging of the presynaptic ATP reporter, Syn-ATP, was performed as previously described (Rangaraju et al., 2014) with the following modifications: 1) all data acquisition and microscope control was carried out remotely, allowing the microscope to be kept in complete light isolation; 2) all images were acquired through a 40x 1.3 NA Fluar Zeiss objective using an ET570LP emission dichroic filter (Chroma) (for fluorescence and luminescence). 3) mCherry fluorescence was excited using a 561nm OBIS laser (Coherent) that was gated to be on only during mCherry image acquisition to avoid creating spurious background light during the luminescence image acquisition; 4) long time series of imaging pairs (fluorescence, luminescence) was automated using a custom-written Andor Basic program; 5) given that we did not observe significant differences in pH changes (Figure S3) when comparing wild type neurons in control, in bafilomycin, and vGlut1-KD neurons, we did not correct ATP measurements for changes in cytosolic pH. ATP_presyn_ is reported as normalized values in each case to the starting luminescence to fluorescence ratio (L/F).

Ouabain [1mM] and bafilomycin [1μM] experiments were done separately with their own paired controls. ATP_presyn_ consumption rate was calculated as the slope of a line fit on L/F control traces between 1 min to 6 min 2DG interval. Our measurement region of interest at each nerve terminal was a circular region of radius ~1 um, corresponding to a volume of ~ 4.2 μm^3^. We previously showed that the average resting ATP_presyn_ concentration is ~1.4mM, which converts to 3.5 x 10^6^ ATP molecules in our measurement volume. The initial rate of decay during 2DG perfusion was 5.4% /min (Figure 1B) corresponding to an initial ATP consumption rate of ~3100 ATP_presyn_ molecules/s.

### pHluorin measurements

pHluorin signals during electrical stimulation are reported as a percentage of the total vesicle pool, whose fluorescence is obtained by perfusion of a Tyrode’s solution containing 50mM NH_4_Cl buffered at pH 7.4 using 25mM HEPES and denoted as ΔF_NH4Cl_ (Sankaranarayanan et al., 2000). pHluorin signals in resting neurons are reported as a percentage of the cytosolic pH = 6.9, obtained as follows:

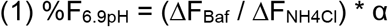

Where α is the correction factor to convert fluorescence determined at pH 7.4 (our NH_4_Cl pH value) to that expected for the pH in the cytosol (pH ~ 6.9) calculated as:

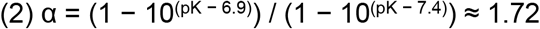

pHluorin pK is 7.1 (Sankaranarayanan et al., 2000).

Images from pHluorin signals in resting neurons were acquired in control Tyrode’s for at least 2min, followed by 500nM bafilomycin perfusion for at least 15min. For experiments inhibiting the vMAT2 transporter, 100nM reserpine was perfused during the entire experiment (with control Tyrode’s and with bafilomycin Tyrode’s).

In experiments using Syphy-pH, Glutamatergic neurons were identified by loading Oyster-550-labeled rabbit anti-vGAT (Synaptic Systems Cat# 131-103C3), as previously described (Pan et al., 2015). Non-labeled boutons were assumed to be Glutamatergic.

### Image Analysis and Statistics

Images were analyzed using the ImageJ plugin Time Series Analyzer V3 where 20-30 circular regions of interest (ROIs) of radius ~ 1μm corresponding to synaptic boutons expressing the pHluorin (as determined with NH_4_Cl superfusion) or SynATP (mCherry positive) were selected and the fluorescence was measured over time. Image loading and posterior raw data saving was automatized by using a homemade python code for Fiji. ROIs signals were analyzed using homemade script routines in Igor-pro V6.3.7.2 (Wavemetric, Lake Oswego, OR, USA). Proton efflux rate was calculated as the slope of a line fit on pHluorin traces between 50s to 400s bafilomycin interval. Results of group data analysis are presented as mean ± SEM. When analyzing means, p values are based on Wilcoxon’s rank test. p < 0.05 was considered significant and denoted with a single asterisk, whereas p < 0.01, p < 0.001 and p < 0.0001 are denoted with two, three, and four asterisks, respectively. The n value, indicated in the figure legends for each experiment, represents the number of cells imaged.

## Data and materials availability

All data is available in the main text or the supplementary materials. Plasmids used for transfection are either available through Addgene or by contacting the authors.

## Acknowledgments

We thank the members of the Ryan lab for helpful discussion and comments in the manuscript. Especial thanks to Andrew Nelson for developing a GUI interface for remote imaging capabilities in the SynATP experiments.

## Author contributions

**Camila Pulido:** Conceptualization, Software, Formal Analysis, Investigation, Writing Original Draft, Visualization. **Timothy A. Ryan:** Conceptualization, Writing Original Draft, Supervision, Project administration, Funding acquisition.

## Additional information

**Supplementary Information** is available for this paper.

**Correspondence and requests for materials** should be addressed to Timothy A. Ryan

---

**Figure S1.**
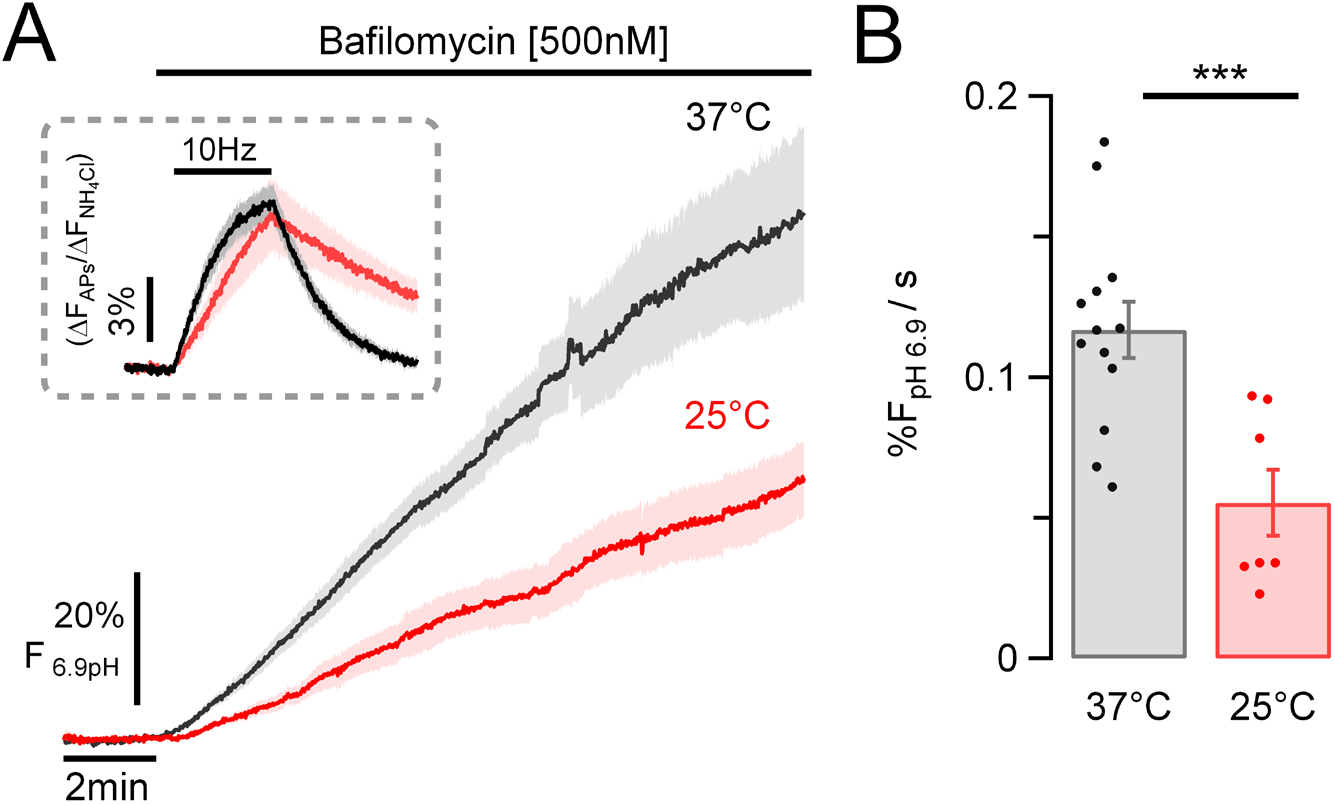
H^+^ efflux is temperature dependent. H^+^ flux from synaptic vesicles measured in hippocampal neurons expressing vG-pH. (A) Average vG-pH traces of control neurons at either 37°C or at 25°C in presence of TTX before and after perfusion with bafilomycin. Fluorescence expressed as a percentage of the fluorescence expected for pH = 6.9 (%F_pH6.9_) based on perfusion with NH_4_Cl (see methods) ± SEM intervals are indicated by shaded colored areas. (Inset) Average vG-pH traces in response to 100APs. (B) Lowering the temperature from 37°C to 25°C reduced the SV H^+^ efflux rate by a factor of ~2. (37°C: 0.117% F_pH6.9_/s ± 0.01 (n = 13) vs 25°C: 0.055% F_pH6.9_/s ± 0.012 (n = 7); p < 0.001)

**Figure S2.**
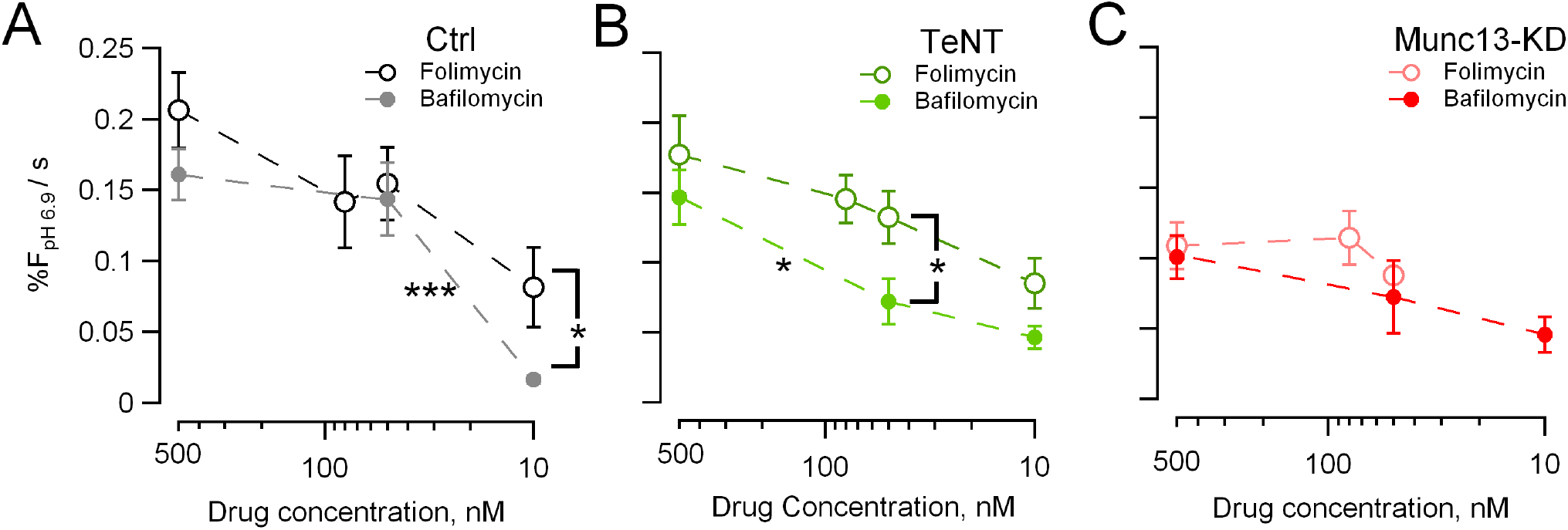
Unmaksing SV H^+^ efflux depends on ATPase block. SV H^+^ efflux from synaptic vesicles expressing vG-pH measured at different concentration of bafilomycin (close symbols) or folimycin (open symbols). Concentration ranged from 10nM – 500nM. (A) Control neurons. In bafilomycin control (gray line, close symbols), H^+^ flux change from 50nM to 500nM was not significant different (50nM: 0.14%F_pH6.9_/s ± 0.025 (n = 8) vs 500nM: 0.16%F_pH6.9_/s ± 0.018 (n = 12); nsd), therefore, we used 500nM as the effective V-ATPase blockage to measure SV H^+^ efflux. (B and C) Neurons where exocytosis was genetically suppressed. (B) Neurons expressing tetanus toxin light chain (TeNT). (C) Neurons suppressing expression of Munc13 (Munc13-KD).

**Figure S3.**
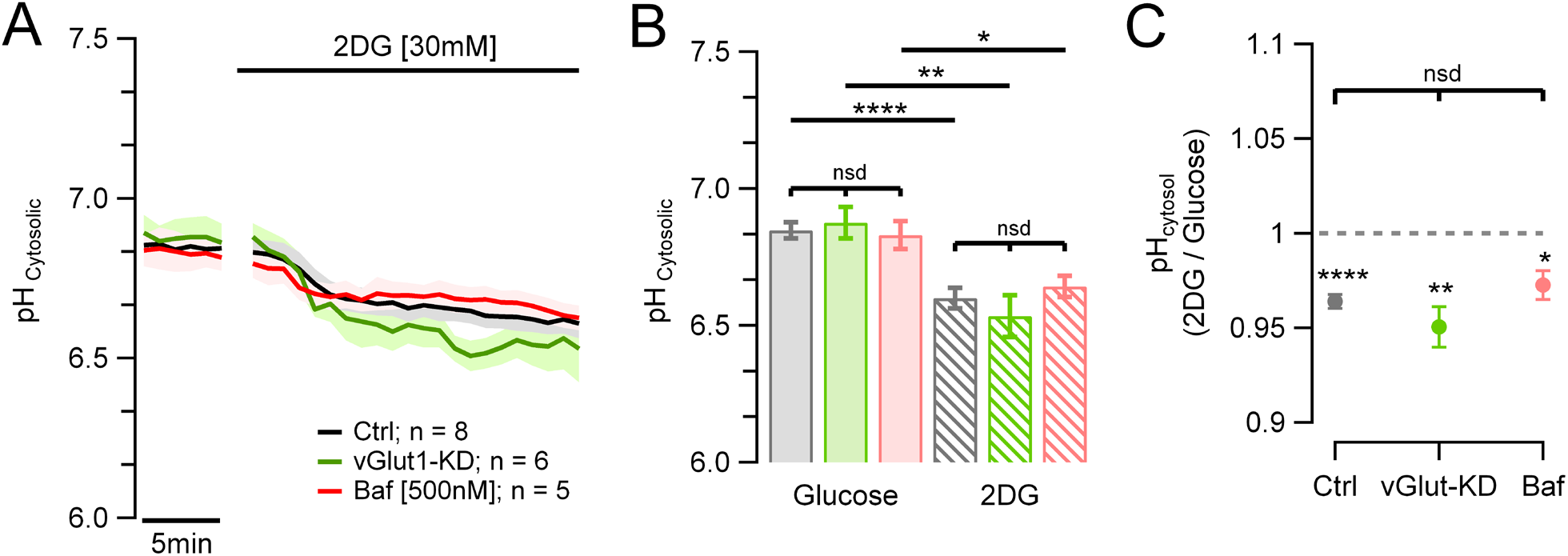
Changes in cytosolic pH during 2DG perfusion is similar in vGlut1-KD neurons or during bafilomycin treatment. Cytosolic pH measurements from hippocampal neurons expressing a cytoplasmic pHluorin. (A) Average cytosolic pH traces in WT neurons in control (n = 8), in Bafilomycin (n = 5) and vGlut1-KD (n = 6) in presence of glucose (first 5min) and 2DG. (B) 2DG causes a cytosolic pH decrease, as previously reported (Rangaraju et al., 2014). Ctrl: pHGlucose 6.85 ± 0.03 vs pH2DG 6.6 ± 0.04 (n = 8; p < 0.0001); bafilomycin: pHGlucose 6.83 ± 0.04 vs pH2DG 6.64 ± 0.04 (n = 5; p < 0.02); vGlut1-KD: pHGlucose 6.87± 0.04 vs pH2DG 6.53 ± 0.08 (n = 6; p < 0.005). (C) Similar pH decrease was observed in control, vGlut1-KD and bafilomycin neurons.

